# *Schmidtea sp*., from the S-W Romania (Platyhelminthes, Tricladida, Dugesiidae)

**DOI:** 10.1101/2021.05.21.445127

**Authors:** Babalean Anda Felicia

## Abstract

The morphology and the anatomy of the copulatory apparatus in a Dugesiidae population from the SW Romania are presented. The copulatory apparatus is characterized by intermingled bursal canal musculature and two distinct penis bulbs with two large seminal vesicles. Based on these morphological characters, the population is assigned to the “*lugubris-polychroa*” group of species, now belonging to the genus *Schmidtea* (de Vries & Sluys 1991). The copulatory apparatus is also characterised by the presence of an atrial fold, characteristic of *S. mediterranea*. The assign of the morphotype here presented to the species level is delayed until integrative molecular analysis.

## Introduction

*Schmidtea* is a genus of freshwater flatworms belonging to Order Tricladida, Suborder Continenticola, Family Dugesiidae (Sluys et al. 2009). It consists of only four species with west-palearctic distribution: *S. lugubris* (Schmidt, 1861), *S. polychroa* (Schmidt, 1861), *S. mediterranea* (Benazzi, Baguñà, Ballester, Puccinelli, del Papa, 1975) and *S. nova* (Benazzi, 1982) (Leria et al. 2018).

In Romania, the genus *Schmidtea* is represented only by the species *lugubris* and *nova. S. nova* was found in the north-western part of Romania at Huedin (Leria et al.2018). The literature registers the presence of *S. lugubris* in the south and east regions - South Plain (Romanian Plain), Dobrogea, Danube Delta, Siret Plain (Năstăsescu 1973) and in the north-western part at Aiud (Leria et al. 2018). On *S. lugubris* for Wallachian Plain (SW Plain or Oltenia Plain), data on the geographical distribution, habitat and biocenoses are brought by Rogoz (1979), yet no anatomical character was published.

The contribution of this paper is the description of the copulatory apparatus in a Schmidtea species from SW Romania.

## Material and methods

Worms were collected by author from ANIF Sadova (National Agency for Land Improvement, 43º 53′ 59.89″ N, 23º 57′ 12.50″ E), with a paintbrush, from the underside of immersed pebbles.

The specimens were fixed in Beauchamp’s fluid (ethanol 96º: 6 volumes, formaldehyde 37%: 3 volumes, glacial acetic acid: 1 volume) for 24 hours, and thereafter stored in 75% ethanol. Histological sections were made at intervals of 5μm and were stained in Haematoxylin-Eosin.

### Abbreviations used in the figures

af: atrial fold, b1: the first penis bulb, b2: the second penis bulb, bc: bursal canal, cb: copulatory bursa, ef: external fold, go: gonopore, ma: male atrium, od: oviduct, pp: penis papilla, sd-v: spermiducal vesicles, sv1: the first seminal vesicle, sv2: the second seminal vesicle, vd: vas deferens

## Results

**Order Tricladida Lang, 1884**

**Suborder Continenticola Carranza, Littlewood, Clough, Ruiz-Trillo, Baguñà & Riutort, 1998**

**Family Dugesiidae Ball, 1974**

**Genus *Schmidtea* Ball, 1974**

### Schmidtea sp

Specimen AFB (nr. 14), collected on 11 July 2017 from the underside of immersed pebbles (Fig. 1a), water temperature: 10º C; air temperature: 37º C; frontal sections on 8 slides,

**Fig. 1:**
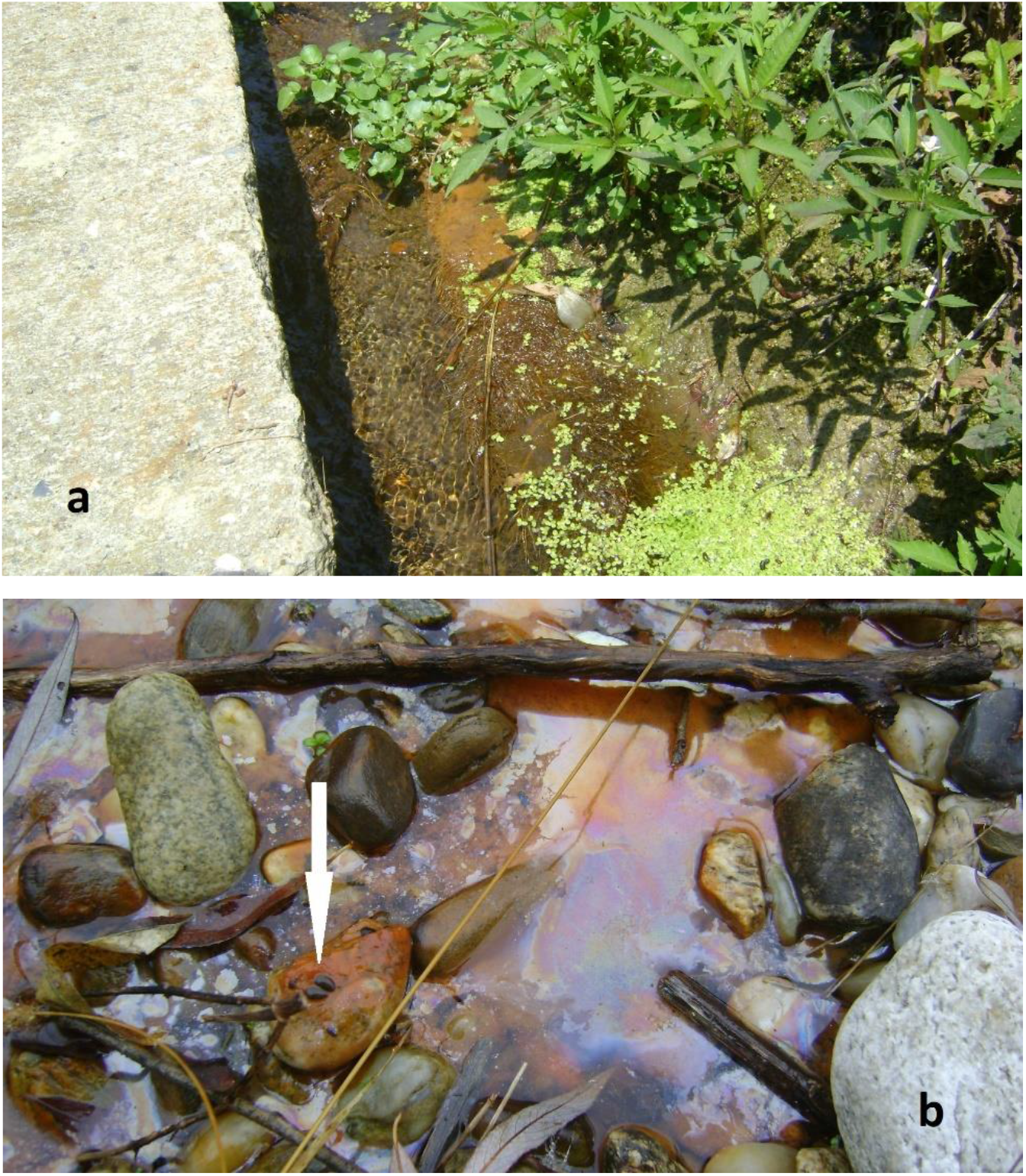
The sampling stations at ANIF Sadova

Specimen AFB (Dg1), collected on 24 November 2017 from the underside of immersed pebbles in a small basin of stagnant water probably polluted by hydrocarbons, judging by the oily appearance of the water surface (Fig. 1b); water temperature: 12º C; air temperature: 10º C; sagittal sections on 32 slides,

Specimen AFB (Dg2), ibid. AFB (DG1), frontal sections on 8 slides.

#### 1. External morphology (Fig. 2)

The size of the sexuate living specimens ranged between 7 mm long (nr. 14, July) to 12 mm long (Dg1, November). The head has a rounded-subtriangular anterior margin and two close eyes set in pigment-free patches. The head narrows very slightly in a neck region. The dorsal surface of the living worm is dark brown; the ventral surface is paler. The fixed worms have a dark brown body with a sandy, granular appearance.

#### 2. Anatomy

The testes (Fig. 3) have a dorsal position in the mesenchyme. On the left and right laterals of the body the testes extend throughout the body length, while in the central part of the body they are more numerous in the pre-pharyngeal region.

**Fig. 2:**
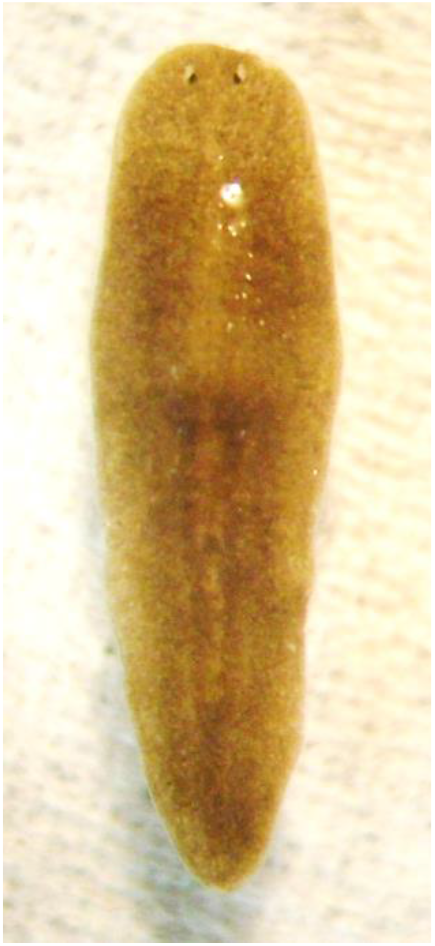
External morphology of a fixed worm (scale bar not available)

**Fig. 3:**
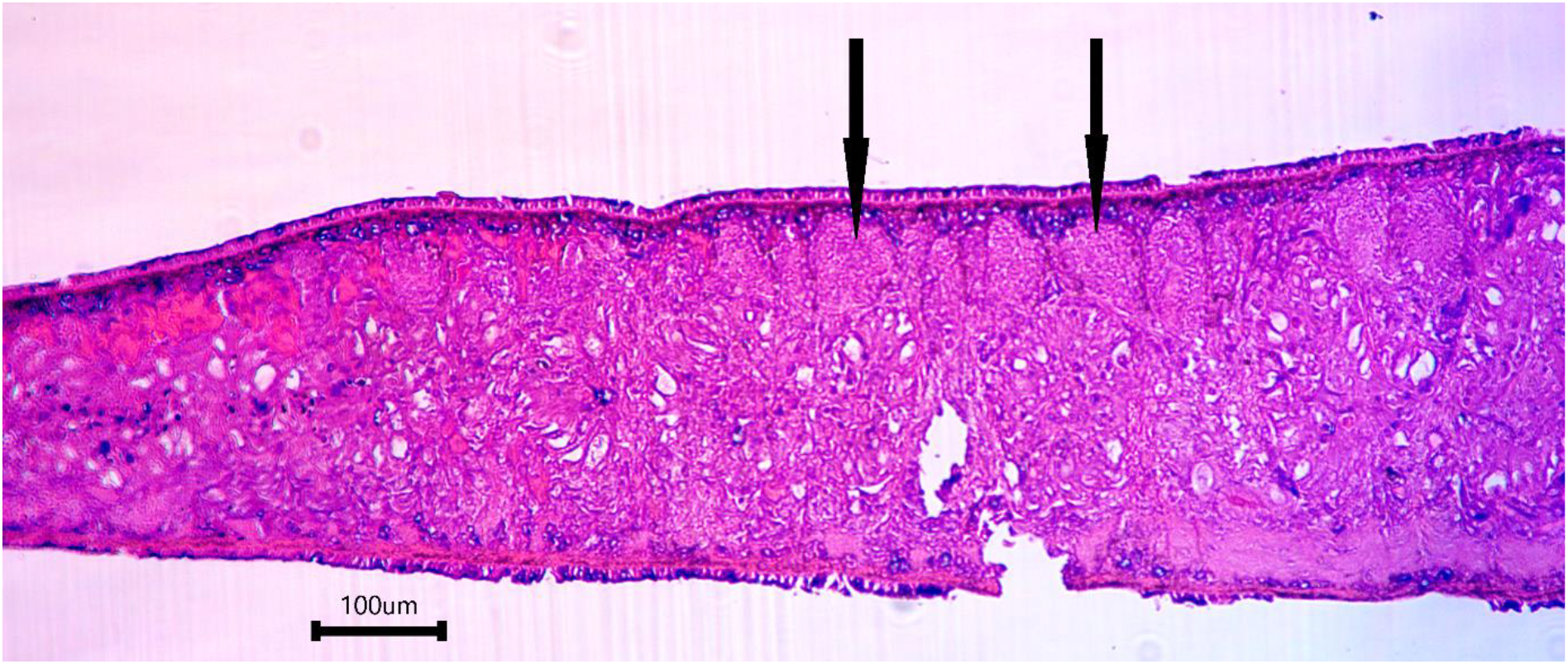
Testes in specimen Dg1 (sagittal section) dorsal testes indicated by arrows

The vasa deferentia (spermiducts) form large spermiducal vesicles on either side of the pharynx. The spermiducal vesicles pass down the level of both bulbs (Fig. 4). Judging by the position of the spermiducal vesicles (Fig. 6), the spermiducts probably take a bent course before opening separately in the posterior wall of the first bulb.

**Fig. 4:**
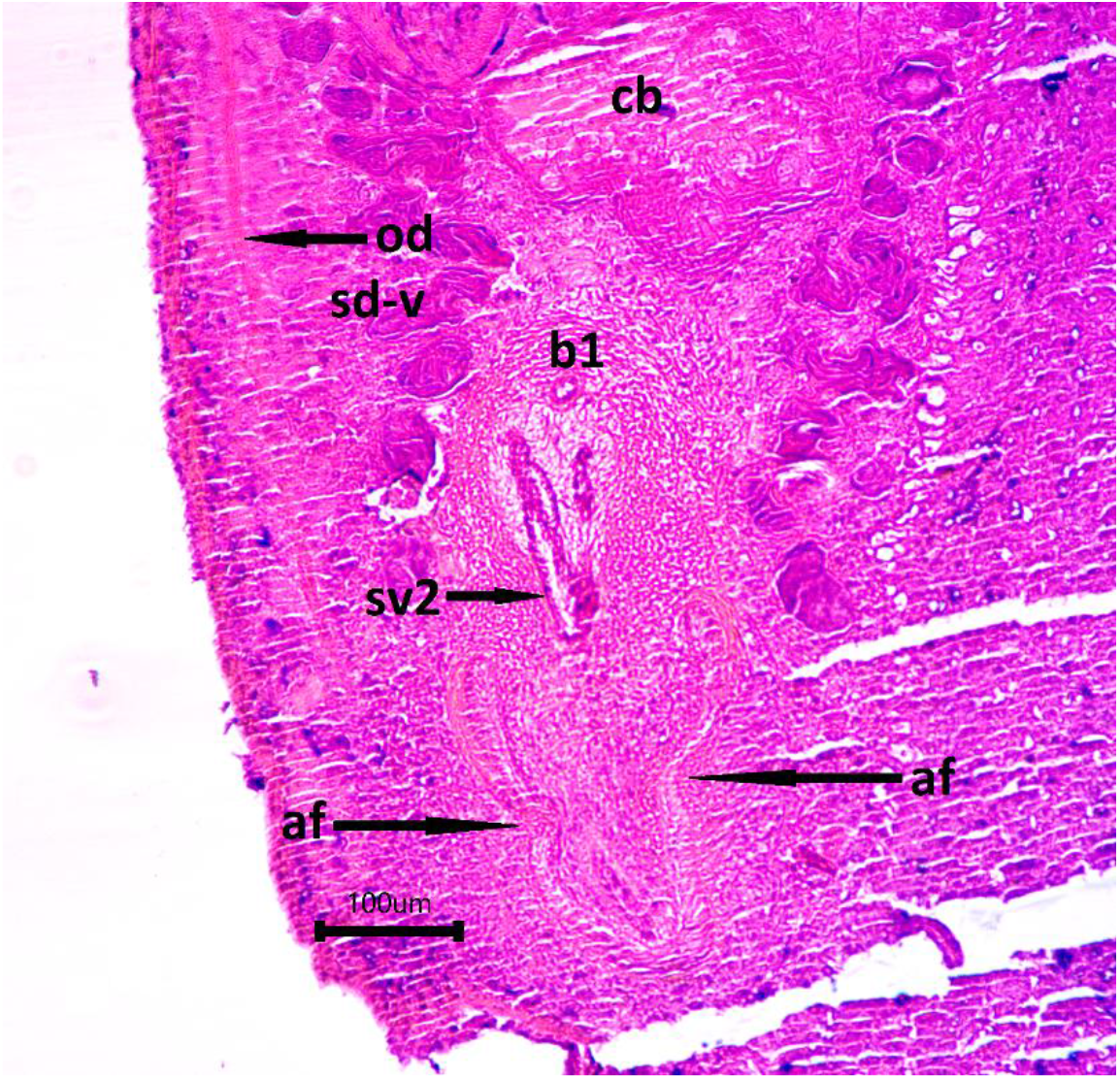
Microphotograph of the copulatory apparatus in specimen Dg2 (frontal section)

The penis (Figs. 4, 5, 6) has two distinct bulbs nearly of the same size: the first bulb (b1) antero-dorsal and the second bulb (b2) postero-ventral. The first bulb presents a developed muscular wall with layers of intermingled and circular musculature. The first seminal vesicle is large and rounded, lined with a wave-shaped epithelium, with cells of different heights. The wall of the second bulb consists of un-oriented musculature. The second seminal vesicle is large and elongated, lined with a tall epithelium with very thin cells. The separation of the penis bulb in two distinct parts is almost imperceptible in the specimen nr. 14.

**Fig. 5:**
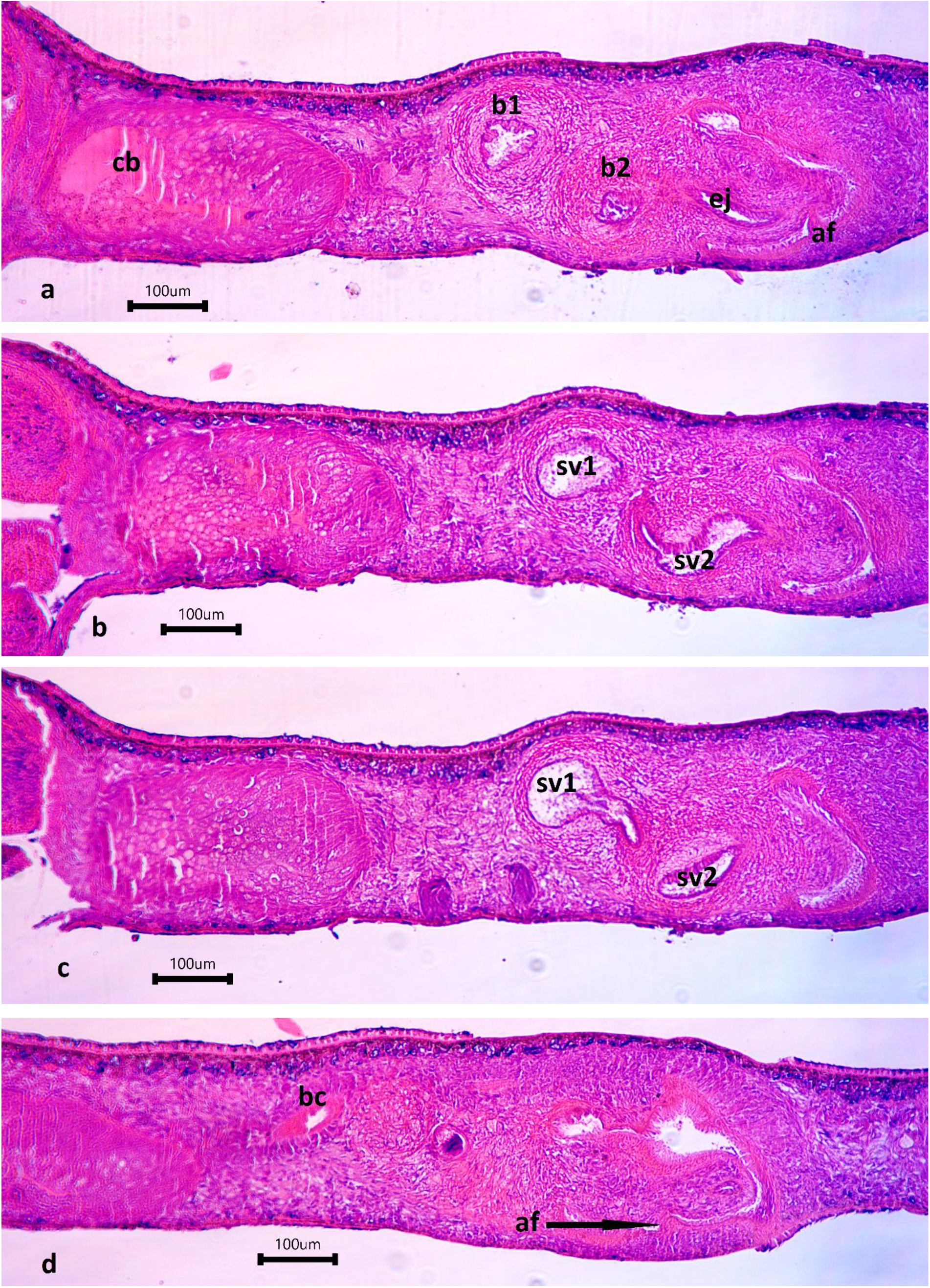
Microphotographs of the copulatory apparatus in specimen Dg1 (sagittal sections)

**Fig. 6:**
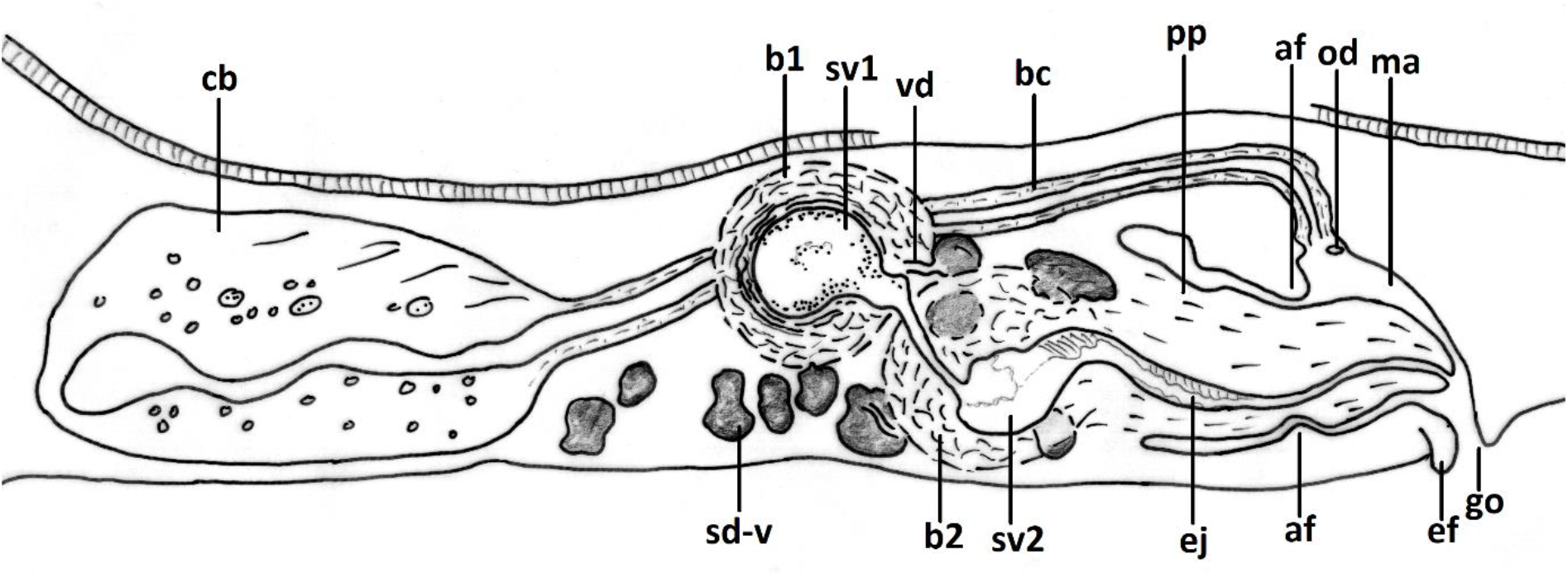
Reconstructed copulatory apparatus in specimen Dg1 af – atrial fold, b1 – the first bulb, b2 – the second bulb, bc – bursal canal, cb – copulatory bursa, ef – external fold, go – gonopore, ma – male atrium, od – oviduct, pp – penis papilla, sd-v – spermiducal vesicle, sv1 – the first seminal vesicle, sv2 – the second seminal vesicle, vd – vas deferens; anterior to the left (scale bar not available)

The two bulbs communicate through a narrow duct (Figs. 5b, c, 6).

The penis papilla is elongate, with a median compression and the ejaculatory duct running slightly ventrally (Fig. 5a). No nipple was observed to the opening of the ejaculatory duct. The ejaculatory duct is lined with a low epithelium sitting on a thin muscular layer.

The epithelium of the genital atrium produces a large fold which compress the penis papilla at it middle (Figs. 4, 5a, d, 6). The genital atrium epithelium reflects outside the gonopore in specimen Dg.1, producing a small external fold. The atrial epithelium consists of low cylindrical, pillar-shape cells, sitting on a thin muscular layer.

The copulatory bursa (Figs. 4, 5, 6) is a large sac lined with a tall, pseudostratified nucleated and vacuolated epithelium covered by a very thin muscular layer; the bursal canal is characterised by intermingled musculature and is lined with a low epithelium, discharging small vacuoles into the lumen.

The two oviducts open separately into the bursal canal, close to where it opens into the male atrium.

## Comparative discussions

The literature presents distinctive morphological characters for each of the four species of the *lugubris-polychroa* complex. For *S. lugubris*, distinctive is considered a permanent nipple on the tip of the penis (Ball & Reynoldson 1981, Reynoldson & Belamy 1970, Leria et al. 2018). *S. polychroa* has particular a dorsal hump on the penis papilla, (Ball & Reynoldson 1981, Leria et al. 2018), a ventral ejaculatory duct and the lack of a permanent nipple (Reynoldson & Belamy 1970). *S. mediterranea* presents a developed atrial fold (Benazzi et al. 1975, Leria et al. 2018). In this four-species group, *S. nova* stands apart by several unique features (Leria et al. 2018).

However, a morphological discrimination between *lugubris, polychroa* and *mediterranea* is not very sharp (precise). For instance, and an atrial fold is also recorded on a small percentage in *S. polychroa* and *S. lugubris* (Benazzi et al. 1975).

Nor does the morphology of the penis bulb appear to be a sharp discriminant character. A penis bulb consisting of two distinct, separated bulbs with two large seminal vesicles nearly of the same size is present in *S. mediterranea* from Italy and Spain (Benazzi et al. 1975, fig. 1, 2, 3) and *S. lugubris* from Romania (Năstăsescu 1973, fig. 3, 4, 5, 7, 8). Instead, in *S. lugubris* of the British fauna, the penis bulb is not separated in two distinct bulbs, not even by an external, superficial constriction, while “two vesicular structures within the bulb” are given: a true anterior, larger seminal vesicle and a smaller, posterior, secondary seminal vesicle (Ball & Reynoldson 1981). In *S. mediterranea* from Tunisia the penis bulb is also figured as two distinct bulbs (Harrath et al. 2004, fig. 1), but not separated. A bulbar constriction is reported by Harrath et al. (2012) in *S. polychroa* from the Palearctic section of the African continent. The penis bulb in *S. nova* consists of two parts separated by a constriction.

Considering the above characters, the worms from the SW Romania at ANIF Sadova differ from *S. lugubris* in lacking a nipple on the tip of the penis and are different from *S. polychroa* in lacking a dorsal hump on the penis papilla. They also differ from *S. lugubris* studied by Năstăsescu (1973) in the presence of an atrial fold, thus pointing to *S. mediterranea*. The bent course of the vasa deferentia before opening into the penis bulb is more characteristic of *S. mediterranea* (Harrath et al. 2004, fig. 1) than of *S. lugubris* presented in the descriptive literature. The external fold in specimen Dg1 might be an individual character or an artefact. The unclear separation of the penis bulb in the case of the small specimen collected in July (specimen nr.14) can lead to misidentification as *S. polychroa*.

It is important to be noticed the communication between the 2 penis bulbs through an external duct in the specimens of *S. lugubris* from Sf. Gheorghe and Mangalia (Năstăsescu 1973, fig. 4, fig. 5).

## Conclusions

1. From a morphological point of view, the specimens subject of this paper, show a more pronounced morphological similarity with *S. mediterranea* than with *S. lugubris*.
2. More and more, the description of new Dugesiid species (and not only), species delineation, inferring phylogenetic relationships rely on integrative methods (e. g. Baguñà et al. 1999, Carranza et al. 1996, Harrath et al. 2019, Lázaro et al. 2009, Lázaro et al. 2011, Leria et al. 2020, Sluys et al. 2013, Song et al. 2020, Stocchino et al. 2013, Wang et al. 2021).

It remains the task of integrative studies to assign the morphotype described in this paper to a species. Additional molecular and morphological analysis could determine whether the atrial fold is a character specific to *S. mediterranea* or is present in other species.

## Acknowledgements

I am deeply grateful to Mrs. Carmen Micu for preparing the histological slides.

